# Characterization of the first cultured free-living representative of *Candidatus* Izimaplasma uncovers its unique biology

**DOI:** 10.1101/2020.11.18.388454

**Authors:** Rikuan Zheng, Rui Liu, Yeqi Shan, Ruining Cai, Ge Liu, Chaomin Sun

## Abstract

*Candidatus* Izimaplasma, an intermediate in the reductive evolution from Firmicutes to Mollicutes, was proposed to represent a novel class of free-living wall-less bacteria within the phylum Tenericutes found in deep-sea methane seeps. Unfortunately, the paucity of marine isolates currently available has limited further insights into their physiological and metabolic features as well as ecological roles. Here, we present a detailed description of the phenotypic traits, genomic data and central metabolisms tested in both laboratorial and deep-sea environments of the novel strain zrk13, which allows for the first time the reconstruction of the metabolic potential and lifestyle of a member of the tentatively defined *Candidatus* Izimaplasma. On the basis of the description of strain zrk13, the novel species and genus *Xianfuyuplasma coldseepsis* is proposed. Notably, DNA degradation driven by *X. coldseepsis* zrk13 was detected in both laboratorial and *in situ* conditions, strongly indicating it is indeed a key DNA degrader. Moreover, the putative genes determining degradation broadly distribute in the genomes of other Izimaplasma members. Given extracellular DNA is a particularly crucial phosphorus as well as nitrogen and carbon source for microorganisms in the seafloor, Izimaplasma bacteria are thought to be important contributors to the biogeochemical cycling in the deep ocean.

## Introduction

The phylum Tenericutes is composed of bacteria lacking a peptidoglycan cell wall and consists of bacteria that evolved from the phylum Firmicutes^1–3^. Two distinctive features set Tenericutes apart from the Firmicutes: the inability to synthetize precursors of peptidoglycan and thereby forming a cell wall^4–7^, and extreme reduction of the size of genome (between 530 to 2220 kbp)^8,9^. Tenericutes includes the class Mollicutes^2^, three taxa in provisional class *Candidatus* Izimaplasma ^4^, and several taxa of unclassified status. Tenericutes bacteria have evolved a broad range of lifestyles, including free-living, commensalism and parasitism^2^. Up to date, almost all reported Mollicutes (including five orders Mycoplasmatales, Entomoplasmatales, Haloplasmatales, Acholeplasmatales, and Anaeroplasmatales)^10^ are commensals or obligate parasites of humans, domestic animals, plants and insects^2^. In comparison, some free-living species are found to be associated with inert substrates (e.g. *Candidatus* Izimaplasma)^4^ or animal/plant surfaces (e.g. *Acholeplasma laidlawii*)^11, 12^.

Tenericutes bacteria ubiquitously exist in numerous environments^13^. Environmental 16S rRNA surveys have identified a large number of unknown Tenericutes clades in diverse environments including the deep oceans, introducing the possibility that these Tenericutes may represent free-living microorganisms which conducting a non-host-associated life style^2^. Indeed, free-living *Candidatus* Izimaplasma^4^ and Haloplasma^14^ were reported in a deep-sea cold seep and brine pool, respectively. These deep-sea free-living Tenericutes exhibit metabolic versatility and adaptive flexibility, showing the possibility to isolate more Tenericutes bacteria from the oceans and even other extreme environments.

Among the free-living Tenericutes identified in the marine environments, *Candidatus* Izimaplasma was mainly studied based on the enrichment and corresponding assembled genomes given the absence of pure cultures^4,15^. *Candidatus* Izimaplasma was proposed to represent a novel class of free-living wall-less bacteria within the phylum Tenericutes^4^, and an intermediate in the reductive evolution from Firmicutes to Mollicutes^4^. Members of the Izimaplasma are thought to be heterotrophs engaging primarily in fermentative metabolism^4^, and thereby producing lactate and possibly other small molecules such as acetate or ethanol, which in turn may be utilized by other members of the cold seep^15^. Notably, based on their genomic analysis, *Candidatus* Izimaplasma bacteria are believed to be important DNA degraders^4,15^. Recent studies have shown high concentrations of extracellular DNA in marine sediments from shallow depths down to the abyssal floor^16–18^. Given extracellular DNA is a particularly crucial phosphorus as well as nitrogen and carbon source for microorganisms in the seafloor, *Candidatus* Izimaplasma might be a key contributor to the biogeochemical cycling in the deep ocean. However, the paucity of isolates currently available has greatly limited further insights into the physiological, ecological and evolutional studies of *Candidatus* Izimaplasma.

In this study, we successfully cultivated the first free-living representative of the class *Candidatus* Izimaplasma. Using growth assay and transcriptomic methods, we further found organic nutrient and thiosulfate could significantly promote the growth of the new isolate. Of note, the novel isolate was found to be an effective degrader of extracellular DNA in both laboratorial and real deep-sea conditions. Lastly, its metabolic pathways were reconstructed based on multi-omics results presented in this study and its potential ecological roles were also discussed.

## Results

### Quantification of the abundance of Tenericutes and *Candidatus* Izimaplasma in the deep-sea cold seep sediments

To gain some preliminary understanding of Tenericutes in the deep-sea, operational taxonomic units (OTUs) sequencing was performed to detect the abundance of Tenericutes present in the cold seep sediments at depth intervals of 0-10 cm, 30-50 cm, 90-110 cm, 150-170 cm, 210-230 cm, 230-250 cm from the surface to the deep layer. As previously reported ^2^, the phylum Tenericutes had a very low abundance in the cold seep sediments (Supplementary Fig. 1a). For example, the percentages of Tenericutes only respectively accounted for 0.011%, 0.222% and 0.003% of the whole bacterial domain in the samples ZC1, ZC3 and ZC4 (Supplementary Fig. 1a). We even could not detect any Tenericutes in other three samples. Thus we propose that extreme low abundance might be a major reason limiting the isolation of bacteria from this phylum. To obtain further insights into the Tenericutes in deep-sea sediments, we performed metagenomics sequencing of the sample ZC3 and analyzed all the annotated genes. The results showed that sequences associated with phylum Tenericutes represented 0.250% of all annotated bacterial sequences (Supplementary Fig. 1b), which was consistent with the OTUs result of the ZC3 sample (Supplementary Fig. 1a). Of note, the proportion of genes associated with *Candidatus* Izimaplasma to those related to Tenericutes was up to 93.289% (Supplementary Fig. 1c), indicating the dominant status of *Candidatus* Izimaplasma within the deep-sea Tenericutes.

### DNA degradation-driven isolation of the first cultured free-living representative of *Candidatus* Izimaplasma from the deep-sea cold seep

Despite what has been learned from cultivation independent methods^4^, the lack of cultured free-living representatives of deep-sea *Candidatus* Izimaplasma has hampered a more detailed exploration of the group. With this, we need cultivated representatives to provide further understanding of *Candidatus* Izimaplasma from the deep-sea environment. In service of this goal, we improved the enrichment strategy by using a specific medium containing only some inorganic salts supplemented with *Escherichia coli* genomic DNA as the nutrient sources, given the previously predicted superior DNA degradation capability of *Candidatus* Izimaplasma^4,15^. Using this medium, we anaerobically enriched the deep-sea sediment samples at 28 °C for six months. Enriched samples were then plated on solid medium in Hungate tubes and individual colonies with distinct morphology were picked and cultured (Fig. 1a). Excitedly, most cultured colonies were identified as members of *Candidatus* Izimaplasma. Among them, strain zrk13 possessed a fast growth rate and was chosen for further study. Under transmission electron microscopy (TEM) observation, the cells of strain zrk13 were coccoid (Figs. 1b and 1e), approximately 300-800 nm in size, and had no flagellum. As expected, cells of strain zrk13 had no distinct cell wall compared with the Gram-positive bacteria cell wall found within members of the Firmicutes (Figs. 1c and 1d), which was confirmed by the TEM observation of ultrathin-sections of strain zrk13 and a typical Firmicutes bacterium (Figs. 1f and 1g). Overall, we successfully obtained the first cultured free-living representative of *Candidatus* Izimaplasma from the cold seep through a DNA degradation-driven strategy, which might be useful to enrich and isolate other uncultured candidates of Izimaplasma in the future.

**Fig. 1.**
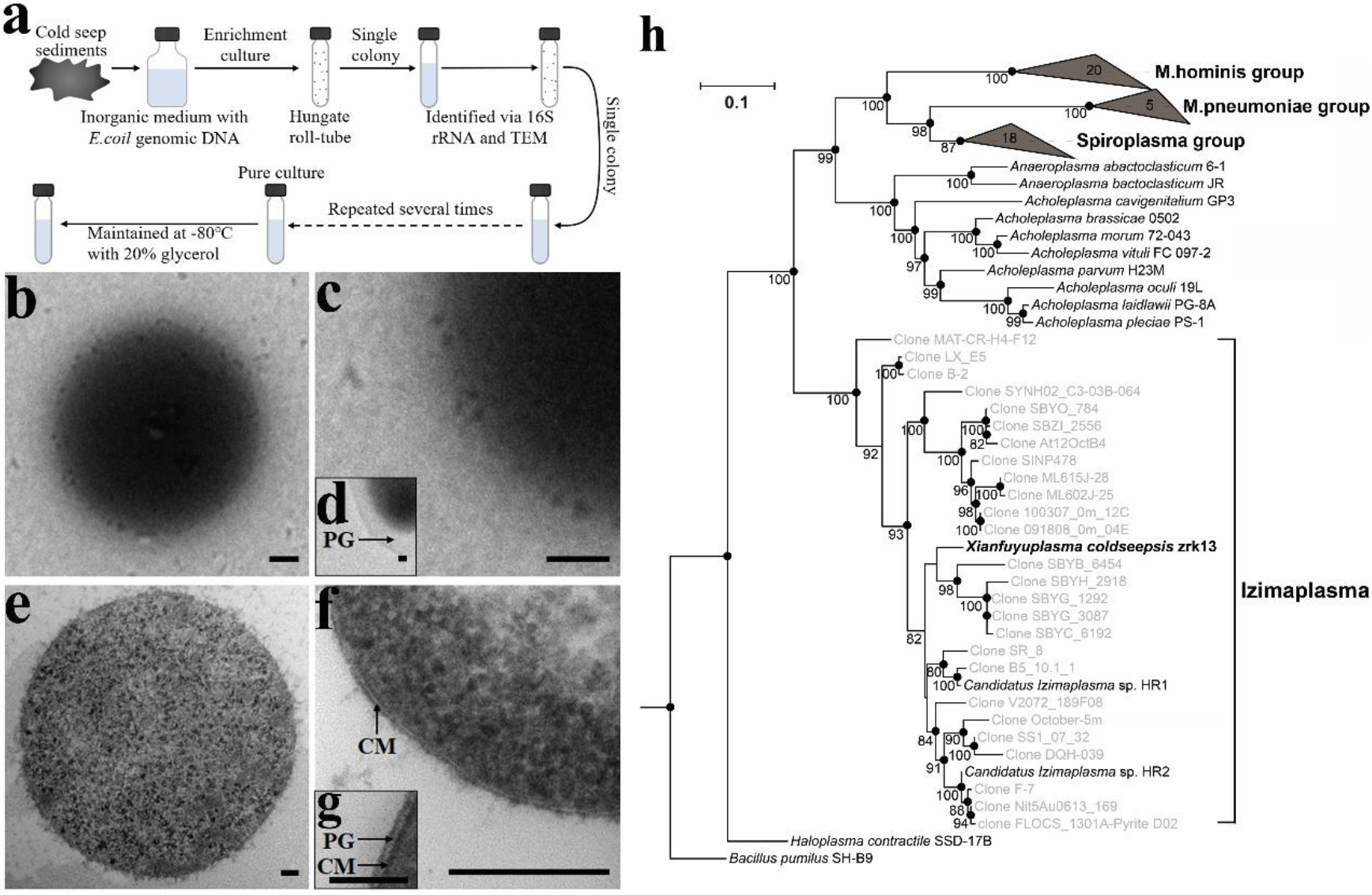
DNA degradation-driven isolation strategy and morphology of *Xianfuyuplasma coldseepsis* zrk13. **a**, Diagrammatic scheme of enrichment and isolation of Izimaplasma bacteria. **b-d**, TEM observation of strain zrk13. **e**, **f**, TEM observation of the ultrathin sections of strain zrk13. **g**, TEM observation of the ultrathin sections of a typical Gram-positive bacterium *Clostridium* sp. zrk8. CM, cell membrane; PG, peptidoglycan; Scale bars, 50 nm. **h**, Phylogenetic analysis of *Xianfuyuplasma coldseepsis* zrk13. Phylogenetic placement of strain zrk13 within the Tenericutes phylum based on almost complete 16S rRNA gene sequences. The tree is inferred and reconstructed under the maximum likelihood criterion and bootstrap values (%) > 80 are indicated at the base of each node (expressed as percentages of 1,000 replications). The black dots at a distinct node indicate that the tree was also formed using the neighbor-joining and minimum-evolution methods. Grey names in quotation represent taxa that are not yet validly published. The only two clades (Mollicutes and Izimaplasma) of Tenericutes are indicated on the right. The sequence of *Bacillus pumilus* SH-B9^T^ is used as an outgroup. Bar, 0.1 substitutions per nucleotide position.

### Genomic characteristics and phylogenetic analysis of strain zrk13

To understand more characteristics of the strain zrk13, its whole genome was sequenced and analyzed. The genome size of strain zrk13 is 1,958,905 bp with a DNA G+C content of 38.2% (Supplementary Fig. 2 and Supplementary Table 1). Annotation of the genome of strain zrk13 revealed it consisted of 1,872 predicted genes that included 64 RNA genes (three rRNA genes, 42 tRNA genes and 28 other ncRNAs). The genome relatedness values were calculated by the amino acid identity (AAI), the average nucleotide identity (ANI), *in silico* DNA-DNA similarity (*is*DDH) and the tetranucleotide signatures (Tetra), against the four genomes (strains zrk13, HR1, HR2 and ZiA1) (Supplementary Table 1). The AAI values of zrk13 with HR1, HR2 and ZiA1 were 68.8%, 66.5% and 64.3%, respectively. The average nucleotide identities (ANIb) of zrk13 with HR1, HR2 and ZiA1 were 68.72%, 67.42% and 66.57%, respectively. The average nucleotide identities (ANIm) of zrk13 with HR1, HR2 and ZiA1 were 85.94%, 86.53% and 81.98%, respectively. The tetra values of zrk13 with HR1, HR2 and ZiA1 were 0.85192, 0.77694 and 0.837. Based on digital DNA-DNA hybridization employing the Genome-to-Genome Distance Calculator GGDC, the *in silico* DDH estimates for zrk13 with HR1, HR2 and ZiA1 were 17.50%, 18.50% and 15.80%, respectively. These results together demonstrated the genome of strain zrk13 to be clearly below established ‘cut-off’ values (ANIb: 95%, ANIm: 95%, AAI: 95%, *is*DDH: 70%, Tetra: 0.99) for defining bacterial species, strongly suggesting it represents a novel taxon within *Candidatus* Izimaplasma as currently defined.

To clarify the taxonomic status of strain zrk13, we further performed the phylogenetic analyses with 16S rRNA genes from cultured Tenericutes and Firmicutes representatives, unclassified *Haloplasma contractile* SSD-17B and some uncultured *Candidatus* Izimaplasma. The maximum likelihood tree of 16S rRNA placed the clade ‘Izimaplasma’ as a sister group of the class Mollicutes, which together form a distinct cluster as Tenericutes separating from the phylum Firmicutes and unclassified *Haloplasma contractile* SSD-17B (Fig. 1h). Of note, the position of *Candidatus* Izimaplasma was just located between the phylum Firmicutes and Mollicutes, indicating *Candidatus* Izimaplasma could represent an intermediate in the reductive evolution from Firmicutes to Mollicutes. Given that strain zrk13 is the first cultured free-living representative of a proposed class *Candidatus* Izimaplasma^4^, which is qualified to be named as Izimaplasma from now on. A sequence similarity calculation using the NCBI server indicated that the closest relative of strain zrk13 was uncultivated Izimaplasma HR1 (95.26%). Therefore, we propose that strain zrk13 is classified as the type strain of a novel genus in the class Izimaplasma, for which the name *Xianfuyuplasma coldseepsis* gen. nov., sp. nov. is proposed.

### Description of *Xianfuyuplasma* gen. nov. and *Xianfuyuplasma coldseepsis* sp. nov

For the genus name *Xianfuyuplasma* Xian.fu.yu’plas.ma. L. fem. n. *Xianfuyu* comes from a strange animal’s name described in the Classic of Mountains and Rivers-a very famous Chinese mythology. *Xianfuyu* is a kind of strange animal possessing a fish’s head and pig’s body (Supplementary Fig. 3), which is similar to Izimaplasma that possessing both characteristics of Tenericutes and Firmicutes. -plasma, formed or molded, refers to the lack of cell wall. The genus *Xianfuyuplasma* is a kind of strictly anaerobic microorganisms, whose cells are non-motile and coccoid. Its phylogenetic position is classified in the family Izimaplasmaceae, order Izimaplasmales, class Izimaplasma within the bacterial phylum Tenericutes. The type species is *Xianfuyuplasma coldseepsis*.

*Xianfuyuplasma coldseepsis* (cold.seep’sis. L. gen. pl. n. *coldseepsis* of or belonging to the deep-sea). For this species, cells are coccoid, approximately 300-800 nm in size, and had no flagellum; strictly anaerobic; the temperature range for growth is 24-32 °C with an optimum at 28 °C. Growing at pH values of 6.0-8.0 (optimum, pH 7.0); growth occurs at NaCl concentrations between 0.0-8.0% with an optimum growth at 4.0% NaCl; growth is stimulated by using glucose, maltose, butyrate, acetate, formate, fructose, sucrose, mannose, starch, isomaltose, trehalose, lactate, ethanol, glycerin and rhamnose as a sole carbon source (Supplementary Table 2), showing quite differences toward the carbon sources predicted by other members of the class Izimaplasma. The type strain, zrk13^T^, was isolated from the sediment of deep-sea cold seep, P. R. China.

### Organic nutrient and thiosulfate significantly promote the growth of *X.coldseepsis* zrk13

In the previous study, uncultivated Izimaplasma HR1 and HR2 were found to be better enriched in the medium amended with small amount of glucose (5 g/L) and yeast extract (0.2 g/L) than that in the inorganic medium^4^. Similarly, the growth rate of *X. coldseepsis* zrk13 was also significantly promoted in the organic medium containing yeast extract (1 g/L) and peptone (1 g/L) compared with that in the oligotrophic medium containing less yeast extract (0.1 g/L) and peptone (0.1 g/L) (Fig. 2a), strongly indicating strain zrk13 is a heterotrophic bacterium. Given the tricarboxylic cycle was incomplete in the previous^4^ and the present Izimaplasma genomes, we sought to ask how does Izimaplasma utilize organic nutrient. Therefore, we performed the transcriptomic analysis of *X. coldseepsis* zrk13 cultured in oligotrophic and rich media. The results revealed that the expressions of genes encoding different glycoside hydrolases (Fig. 2b), FeFe hydrogenases (Fig. 2c), NADH-ubiquinone oxidoreductase (Fig. 2d) and ATP synthase (Fig. 2e) were significantly up-regulated. Glycoside hydrolases are enzymes that catalyze the hydrolysis of the glycosidic linkage of glycosides, leading to the formation of small molecules sugars for easy utilization by organisms (http://www.cazypedia.org/index.php/Glycoside_hydrolases). Correspondingly, expressions of different glycoside hydrolases including families 1, 16, 17, 30, 35, 65 were evidently up-regulated, suggesting many categories of glycans could be efficiently degraded by *X. coldseepsis* zrk13. Indeed, all the present and two uncultured Izimaplasma HR1 and HR2 genomes contained a complete set of genes associated with Embden-Meyerhof-Parnas (EMP) glycolysis and pentose phosphate pathways^4^, indicating Izimaplasa bacteria possess a significant capability of glycan metabolism. Of note, the expressions of all three iron hydrogenase maturation genes *(hydEFG)* were up-regulated about 30-fold (Fig. 2b), and this kind of hydrogenase is usually linked with NADP and may generate NADPH for the production of biomolecules^4^. Correspondingly, the expressions of many genes related to NADH-ubiquinone oxidoreductase were up-regulated 20-fold (Supplementary Fig. 7), which catalyzed the transfer of two electrons from NADH to reduce ubiquinone to ubiquinol and was also the entry point for a large fraction of the electrons that traverse the respiratory chain^19,20^. With this, it is reasonable to see that the expressions of different genes encoding ATP synthase were also greatly up-regulated (Fig. 2e), which converted the energy of protons (H^+^) moving down their concentration gradient into the synthesis of ATP and thereby promoting the bacterial growth. Overall, we conclude that *X. coldseepsis* zrk13 could effectively utilize organic nutrients for growth through anaerobic fermentation and has the potential to produce hydrogen^4^.

**Fig. 2.**
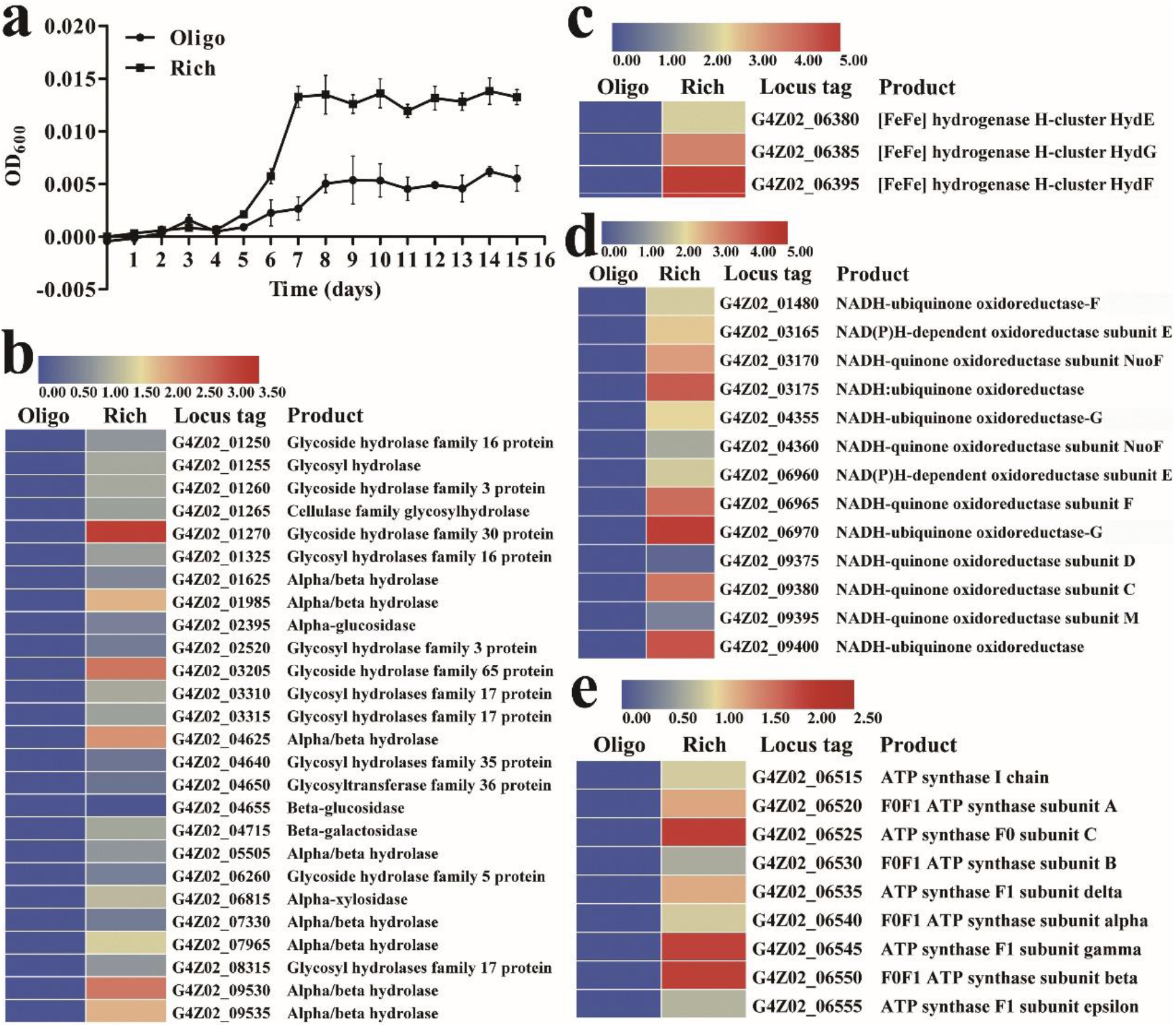
Organic nutrient significantly promotes the grwoth of *X. coldseepsis* zrk13. **a**, Growth assays of strain zrk13 in the oligotrophic and rich media. **b**, Transcriptomics based heat map showing all up-regulated genes encoding glycosyl hydrolases. **c**, Transcriptomics based heat map showing three up-regulated iron hydrogenase maturation genes (*hydEFG*). **d**, Transcriptomics based heat map showing all up-regulated genes encoding NADH-quinone/ubiquinone oxidoreductase. **e**, Transcriptomics based heat map showing all up-regulated genes encoding ATP synthase. Oligo indicates the oligotrophic medium; rich indicates the rich medium.

*X. coldseepsis* zrk13 was isolated from a deep-sea cold seep, and different sulfur sources were detected to ubiquitously exist in the environment in our previous study^21^. Therefore, we tested the effects of different sulfur-containing inorganic substances (e. g. Na_2_SO_4_, Na_2_SO_3_, Na_2_S_2_O_3_, Na_2_S) on the growth of *X. coldseepsis* zrk13. The results showed that only the supplement of Na_2_S_2_O_3_ could significantly promote the growth of strain zrk13 (Fig. 3a). We thus performed the transcriptomic analysis of strain zrk13 cultured in the organic medium amended with or without Na_2_S_2_O_3_ to explore the underlying mechanism of growth promotion. Surprisingly, we didn’t find obvious up-regulation of typical genes associated with sulfur metabolism. Alternatively, the expressions of many genes encoding [2Fe-2S] binding proteins or [4Fe-4S] di-cluster domain-containing proteins were markedly up-regulated (Fig. 3b), and these proteins are thought to play a variety of functions including substrate binding and activation, electron transport, gene expression regulation and enzyme activity in response to external stimuli^22^. Meanwhile, the expressions of large amount of genes encoding enzymes responsible for saccharides degradation (Fig. 3c) and sugar transport (Fig. 3d) were evidently up-regulated. In combination of the results showed in Fig. 2, we speculate that thiosulfate may accelerate the hydrolysis and uptake of saccharides under the control of [Fe-S] associated proteins and thereby synthesizing energy and promoting the growth of strain zrk13. Together, we believe *X. coldseepsis* zrk13 possesses a capability to utilize both organic nutrients (saccharides) and inorganic sulfur-containing compounds (thiosulfate) ubiquitously existing in the deep-sea environments.

**Fig. 3.**
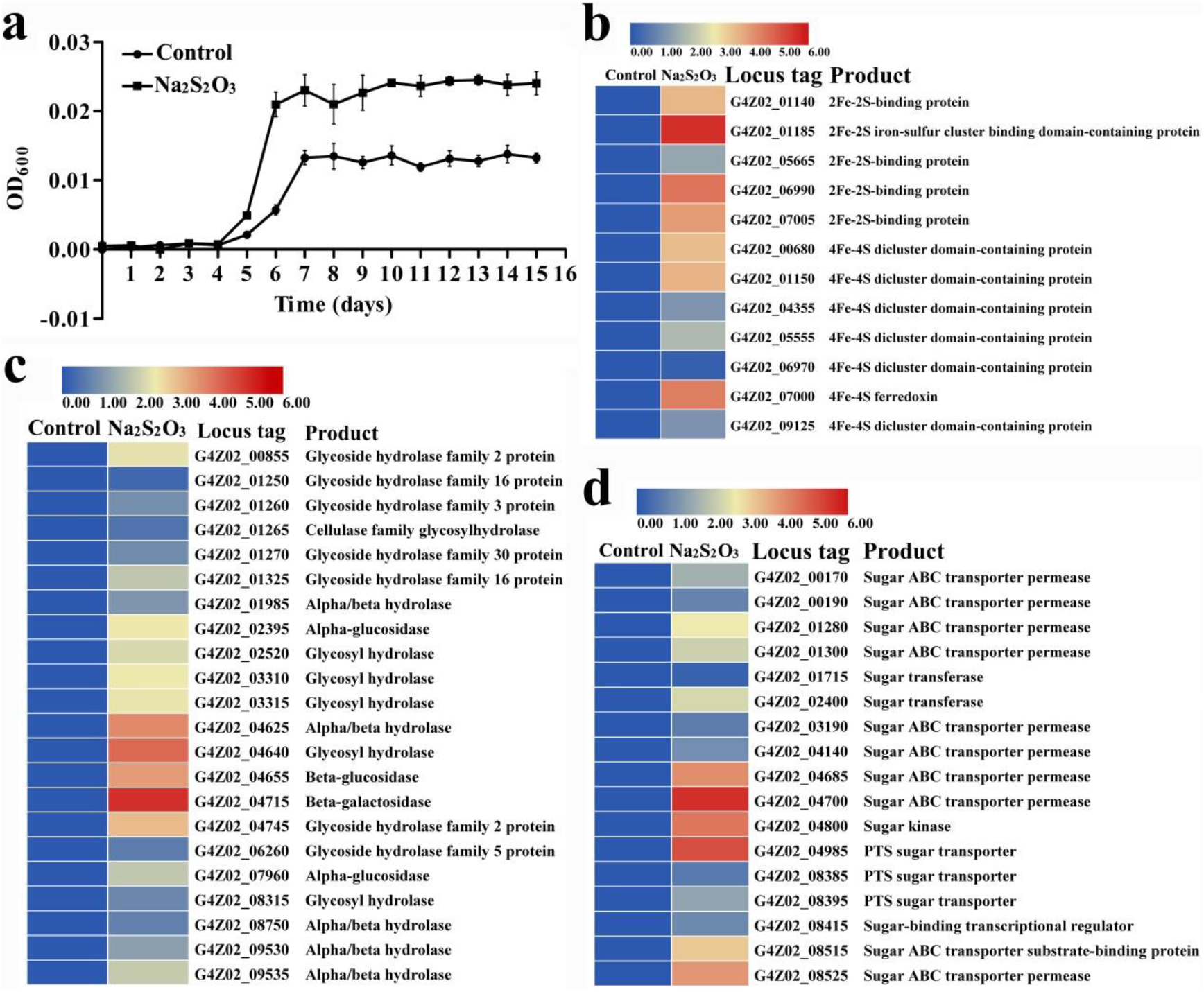
Thiosulfate significantly promotes the grwoth of *X. coldseepsis* zrk13. **a**, Growth assays of strain zrk13 in the organic medium supplemented with or without 100 mM Na_2_S_2_O_3_. **b**, Transcriptomics based heat map showing all up-regulated genes encoding 2Fe-2S/4Fe-4S-binding proteins. **c**, Transcriptomics based heat map showing all up-regulated genes encoding glycosyl hydrolases. **d**, Transcriptomics based heat map showing all up-regulated genes encoding sugar ABC transporter permease.

### *X.coldseepsis* zrk13 possesses a significant capability of degrading DNA in both laboratorial and deep-sea environments

Based on several previous reports^2,4,15^ and our present isolation strategy, Izimaplasma is firmly believed to degrade extracellular DNA and use as a nutrient source for growth. However, this hypothesis is still lack of solid proof due to the uncultivation status of Izimaplasma. With that, detection of DNA degradation and utilization was further performed with strain zrk13. First, an in-depth analysis of the genome of strain zrk13 revealed the existence of a putative DNA-degradation related gene cluster, which contained genes encoding DNA degradation protein EddB, DNase/RNase endonuclease, thermonuclease, phosphohydrolases, ABC-transporters, phosphate transporters and other proteins associated with DNA degradation (Fig. 4a). In order to test the ability of strain zrk13 to degrade DNA, genomic DNA without removing rRNA of *E. coli* was extracted for further assays. The results showed that the supernatant of zrk13 cultures began to digest chromosomal DNA within 5 min (Fig. 4b, lane III) and degraded all DNA within 15 min (Fig. 4b, lane IX). However, the supernatant of zrk13 didn’t degrade RNA at all, indicating its specific DNA degradation capability (Fig. 4b). Moreover, the supplement of genomic DNA in the inorganic medium could significantly promote the growth rate and biomass amount of zrk13 (Fig. 4c), strongly suggesting this bacterium could efficiently utilize the digested DNA as a nutrient source to support further growth. Consistently, the expressions of genes related to DNA degradation were significantly up-regulated, especially the gene encoding DNA degradation protein EddB, which was up-regulated about 20-fold (Fig. 4d), indicating the key roles of the DNA-degradation gene cluster in the course of DNA degradation and utilization. Notably, compared with 62 finished and draft genomes of Tenericutes and Firmicutes bacteria, the members of Izimaplasma harbored more genes associated with DNA degradation than those in Mollicutes and Firmicutes (Supplementary Table 3 and Supplementary Fig. 4), confirming the strong capability of DNA degradation of Izimaplasma.

**Fig. 4.**
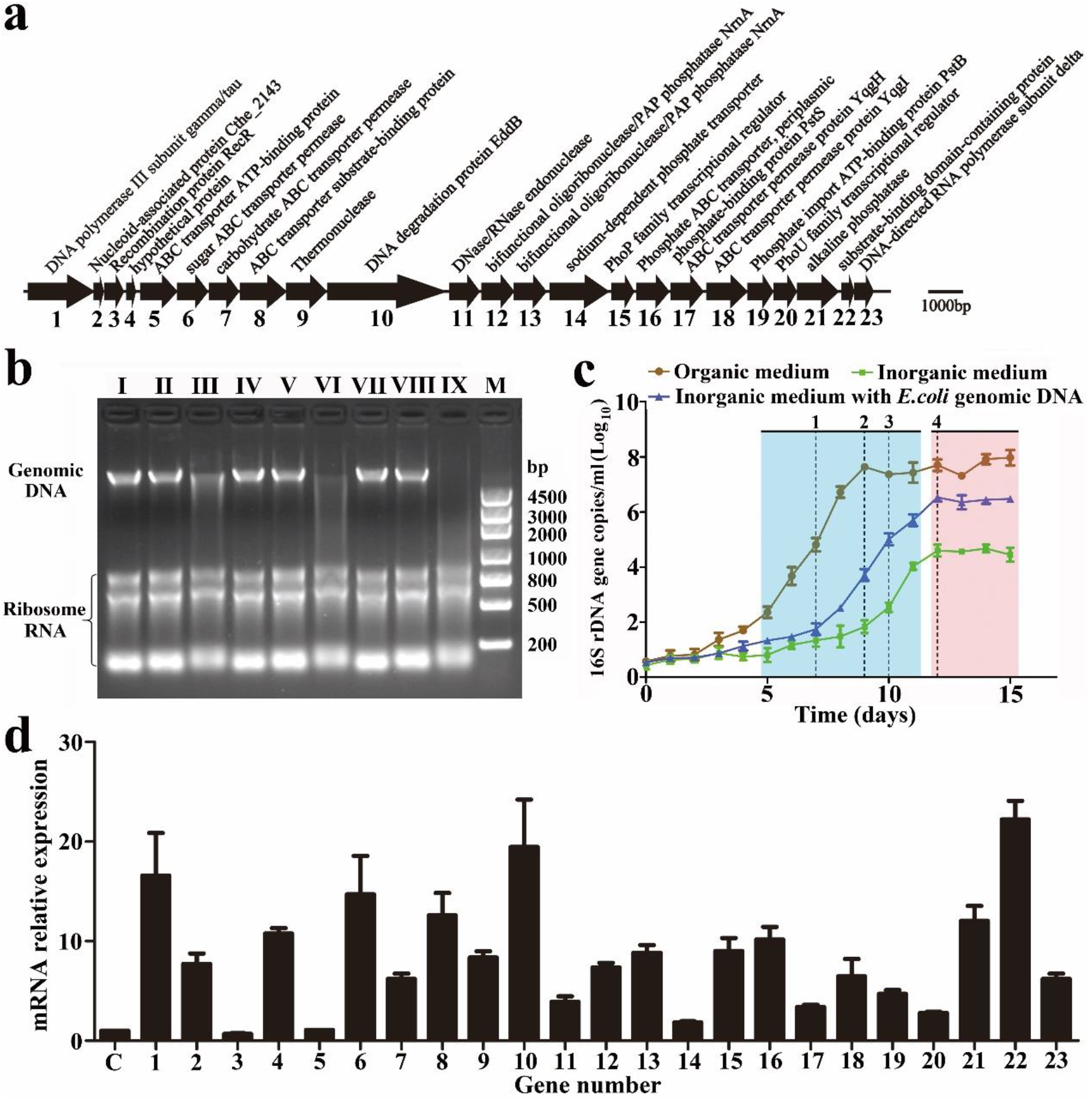
*X. coldseepsis* zrk13 possesses a significant capability of degrading and utilization of extracellular DNA. **a**, Gene arrangements of a putative DNA-degradation locus in strain zrk13. The numbers shown in the X-axis indicate the gene number in the gene cluster. **b**, Detection of the ability of DNA degradation of strain zrk13 by electrophoresis. Lane I, lane IV and lane VII indicate 2 μg *E. coli* genomic DNA and rRNA without treatment. Lane II, lane V and lane VIII indicate 2 μg *E. coli* genomic DNA and rRNA treated by a Clostridia bacterial supernatant for 5 min, 10 min and 15 min at 37 °C, respectively. Lane III, lane VI and lane IX indicate 2 μg *E. coli* genomic DNA and rRNA treated by strain zrk13 supernatant for 5 min, 10 min and 15 min at 37 °C, respectively. M, DL4500 molecular weight DNA marker. **c**, Growth assays of strain zrk13 cultivated in organic medium and inorganic medium supplemented with or without 2 μg *E. coli* genomic DNA. **d**, qRT-PCR detection of expression changes of genes shown in panel a when strain zrk13 cultivated in the inorganic medium supplemented with or without 2 μg *E. coli* genomic DNA. Three biological replicates were performed. The numbers shown in the X-axis indicate the gene number shown in panel a. “C” indicates control.

Taken all the above results, we are clear about the metabolic landscape of *X. coldseepsis* zrk13 based on the laboratorial conditions. However, its real metabolisms performed in the deep-sea environment are still obscure. With that, we performed the *in situ* cultivation in the June of 2020 to study the metabolisms of strain zrk13 in the deep-sea cold seep, where we isolated this bacterium and lives a lot of typical cold seep animals such as mussels and shrimps (Figs. 5a, 5b). The transcriptomics results based on the *in situ* cultivated cells showed that the expressions of many genes encoding exonuclease, endonuclease and ribonuclease were obviously up-regulated compared with control group (Fig. 5c), indicating DNA-degradation indeed happened in the deep sea. Correspondingly, the expressions of all genes belonging to a gene cluster closely associated with purine nucleobase catabolism (such as adenine deaminase, guanine deaminase and xanthine dehydrogenase) were significantly up-regulated up to 16-fold (Fig. 5d), strongly suggesting zrk13 had a remarkable ability to degrade purine and obtain energy to maintain growth *in situ.* Surprisingly, the expressions of almost all the genes involved in EMP glycolysis were significantly down-regulated (Figs. 5e and 5f), which might be due to the deficiency of organic nutrients in the deep-sea environment. Instead, strain zrk13 could be able to maintain a good growth status mainly by digesting DNA to obtain energy for supporting growth given its strong DNA degradation capability and the ubiquitous existence of extracellular DNA in the deep-sea sediments^15,23^.

**Fig. 5.**
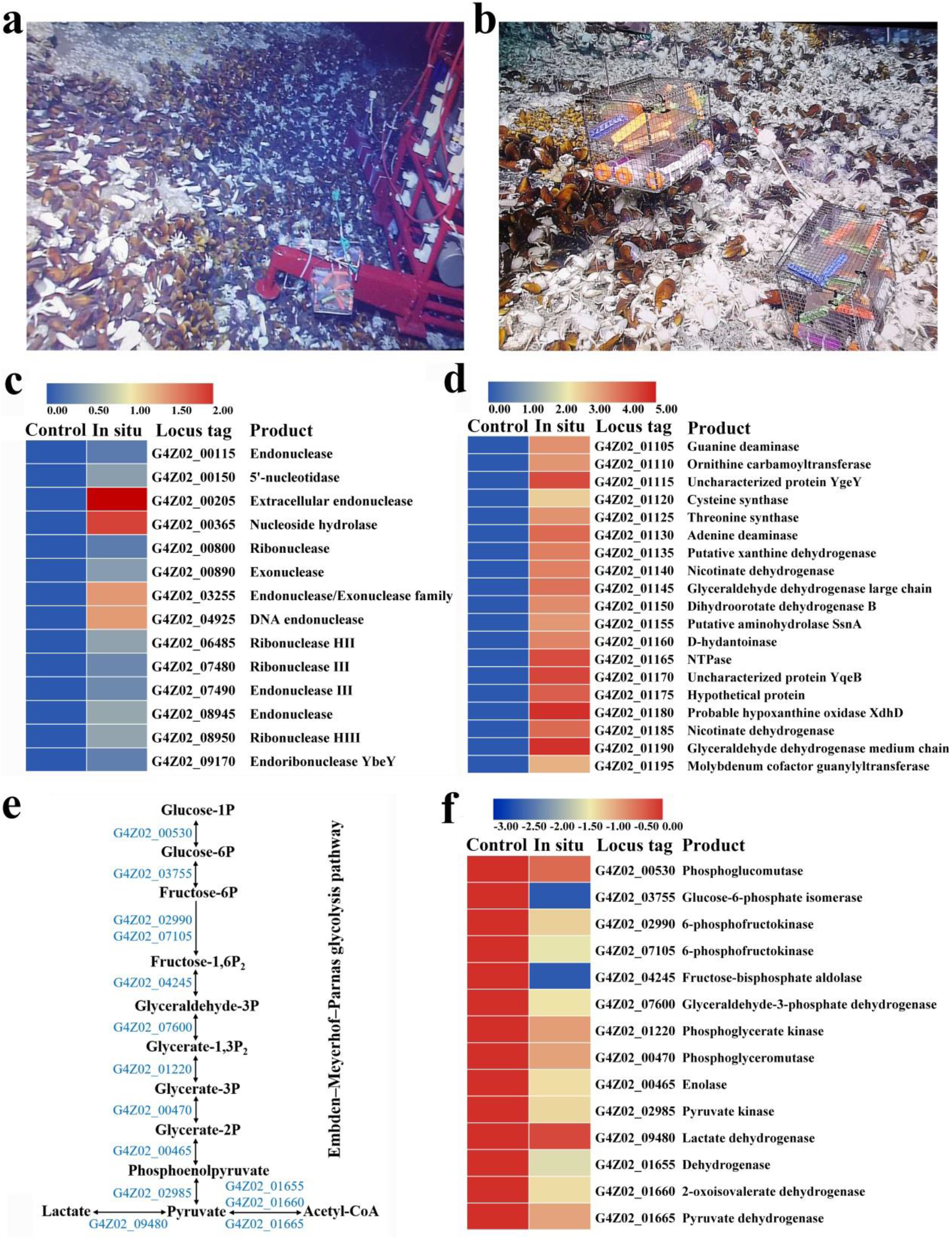
Transcriptomics analysis of *X. coldseepsis* zrk13 incubated in the deep-sea cold seep. **a**, **b**, Distant (a) and close (b) views of the *in situ* experimental apparatus in the deep-sea cold seep where distributed many mussels and shrimps. **c**, Transcriptomics based heat map showing all up-regulated genes encoding enzymes degrading nucleic acids after a 10-day incubation of strain zrk13 in the deep-sea cold seep. **d**, Transcriptomics based heat map showing all up-regulated genes associated with purine catabolism after a 10-day incubation of strain zrk13 in the deep-sea cold seep. **e**, Diagrammatic scheme of EMP glycolysis pathway. The gene numbers showing in this scheme are the same with those shown in panel f. **f**, Transcriptomics based heat map showing all down-regulated genes associated with EMP glycolysis pathway after a 10-day incubation of strain zrk13 in the deep-sea cold seep.

## Discussion

Microbes in deep ocean sediments represent a large portion of the biosphere, and resolving their ecology is crucial for understanding global ocean processes^24–26^. Despite the global importance of these microorganisms, majority of deep-sea microbial diversity remains uncultured and poorly characterized^24^. Description of the metabolisms of these novel taxa is advancing our understanding of their biogeochemical roles, including the coupling of elements and nutrient cycling, in the deep oceans. Given the importance of these communities to the oceans, there is an urgent need to obtain more uncultivated isolates for better resolving the diversity and ecological roles of these uncultured taxa. *Candidatus* Izimaplasma bacteria were proposed to represent a novel class of free-living representatives from a Tenericutes clade found in deep-sea methane seeps^4^, and they were believed to actively participate in the primary degradation of extracellular DNA in anoxic marine sediments and contribute to the biogeochemical cycling of deep biosphere^4,15^. Unfortunately, up to date, there is no any available pure culture of Izimaplasma bacteria, which seriously hinders the accurate determination of their features such as growth, metabolism, physiology and ecology.

In the present study, we developed an effective enrichment method driven by DNA-degradation (Fig. 1a), and successfully obtained a novel Izimaplasma isolate, *X. coldseepsis* zrk13, which is the first pure culture in the Izimaplasma class. Based on the isolate, we detailedly explored its physiology, genomic traits, phylogenetics and metabolisms through bioinformatics, biochemical and transcriptomic methods, and proposed a model describing its central metabolic pathways (Fig. 6). Clearly, DNA and organic matter degradation and utilization play key roles for energy production and thereby driving growth of *X. coldseepsis* zrk13 (Fig. 6).

**Fig. 6.**
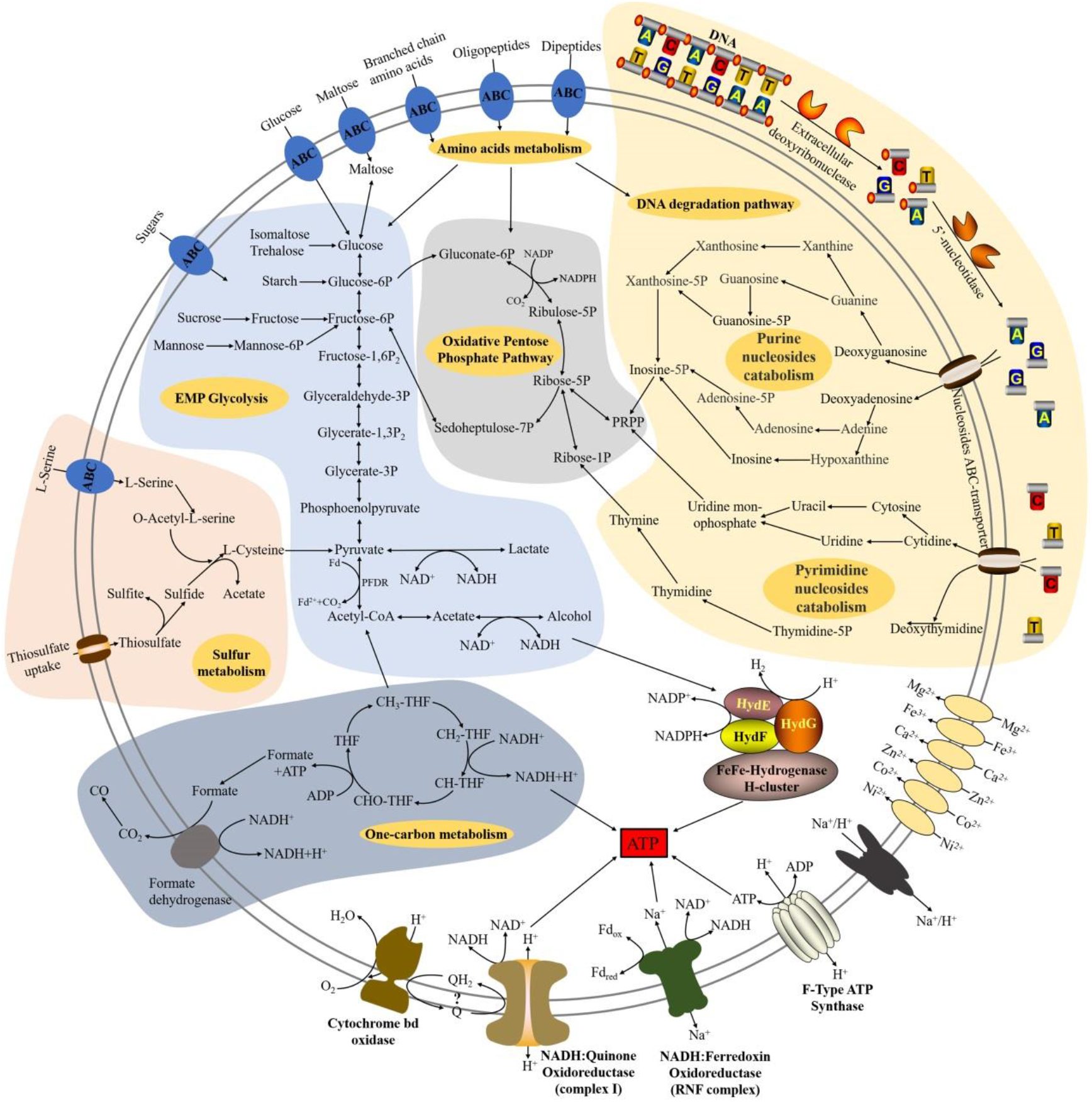
Muti-omics based central metabolisms model of *X. coldseepsis* zrk13. In this model, central metabolisms including DNA degradation, EMP glycolysis, oxidative pentose phosphate pathway, hydrogen production, electron transport system, sulfur metabolism and one carbon metabolism are shown. All the above items are closely related to the energy production in *X. coldseepsis* zrk13. In detail, strain zrk13 contains a complete set of genes related to DNA-degradation, which could catabolically exploit various sub-components of DNA, especially purine-based molecules. And further degradation into nucleotides and nucleosides by nucleases were required to facilitate the introduction of DNA sub-components into cells. Once imported into the cytoplasm, purine- and pyrimidine-deoxyribonucleosides were further broken down into different bases, whereby the respective bases may enter catabolic pathways. The metabolites of nucleosides catabolism were finally transformed into phosphoribosyl pyrophosphate (PRPP) and Ribose-1P, thus entering the oxidative pentose phosphate pathway, which is closely related to EMP glycolysis pathway. Sulfide generated by sulfur reduction from thiosulfate works together with the L-serine to form acetate and L-cysteine, which eventually enter the pyruvate synthesis pathway. Furthermore, the formate dehydrogenase linked with NADH may convert formate into CO2/CO with the production of energy. Iron hydrogenases are linked with NADP and thereby generating NADPH for the production of biomolecules. A membrane-bound, Na^+^-transporting NADH: Ferredoxin oxidoreductase (RNF complex), the H^+^-transporting NADH: Quinone oxidoreductase (complex I) and F-type ATP synthase required for energy metabolism are present in strain zrk13 genome. Both complex I and the cytochrome bd oxidase interact with the quinone pool, which are associated with energy production.

In the laboratorial culturing condition, *X. coldseepsis* zrk13 prefers to perform a heterotrophic life given the easily available organic matter in the medium (Fig. 2a). Therefore, the expressions of large amount of genes associated with sugar degradation were significantly up-regulated compared with the bacterium cultured in the oligotrophic medium (Fig, 2b). In contrast, the expressions of genes responsible for sugar metabolisms (EMP glycolysis) were down-regulated in the *in situ* cultivated *X. coldseepsis* zrk13 (Figs. 5e and 5f) compared with that incubated in the rich medium. Considering the exchanges between the medium inside the incubation bag and the outside seawater, we speculate that the down-regulation of saccharide metabolism *in situ* might be the lower content of organic matters in the seawater than that in the organic medium. Notably, thiosulfate also promotes the saccharides metabolism and thereby benefiting the bacterial growth (Fig. 3), which endows *X. coldseepsis* zrk13 a better adaptability to the deep-sea environment given ubiquitous existence of thiosulfate in the cold seep^21^. Therefore, we believe saccharide metabolism is crucial for maintaining the growth of *X. coldseepsis* zrk13 both in the laboratorial and deep-sea environments. In combination of previous report^4^, we predict part of the ecological role of Izimaplasma in the cold seep is likely in the fermentation of products from the degradation of organic matter to produce lactate and possibly other small molecules such as acetate or ethanol, which in turn are utilized by other members of the cold seep microbes (Fig. 6).

In comparison, the degradation of DNA mediated by *X. coldseepsis* zrk13 happened in both laboratorial (Fig. 4) and deep-sea conditions (Fig. 5), strongly indicating this bacterium is indeed a key DNA degrader. Consistently, an intact locus containing genes encoding enzymes for further digestion of DNA into oligonucleotides and nucleosides, as well as removal of phosphates from nucleotides is identified in the genome of *X. coldseepsis* zrk13 (Fig. 4a). And nucleases within and outside of this gene cluster were demonstrated to function in the course of DNA degradation (Fig. 4 and Fig. 5). Comparative genomics indicated that DNA can be digested by diverse members of Izimaplasma^15^ as well as *X. coldseepsis* zrk13 (Supplementary Fig. 4), strongly suggesting Izimaplasma bacteria are specialized DNA-degraders that encode multiple extracellular nucleases. DNA degradation process is closely associated with amino acids and saccharides metabolisms (Fig. 6) as well as the production of acetyl-CoA^4^. The products of nucleic acids (such as urea and ammonium, CO2 and acetate) are also important nutrients for other members of microbial communities^4,15^.

It is noting that extracellular DNA is a major macromolecule in global element cycles, and is a particularly crucial source of phosphorus, nitrogen and carbon for microorganisms in the seafloor^15^. It was estimated that extracellular DNA available for degradation by extracellular nucleases, supplies microbial communities of both coastal and deep-sea sediments with 2-4% of their carbon requirements, 4-7% of their nitrogen needs, and a remarkable 20-47% of their phosphorus demands^23,27^. Considering the tremendous body of marine water, the oceans harbor a massive Izimaplasma population composed of diverse novel linages. Therefore, extracellular available DNAs and their key degraders like Izimaplasma contribute substantially to oceanic and sedimentary biogeochemical cycles, and provide an enormous energy source for microbial communities. Overall, based on the comprehensive investigations of the first cultured isolate of Izimaplasma in both laboratorial and deep-sea conditions, our present study expands the ecophysiological understanding of Izimaplasma bacteria by showing that they actively participate in the primary degradation of extracellular DNA and other organic matter in anoxic deep-sea sediments.

## Methods

### Samples sampling and statistical analysis of 16S rRNA genes of Tenericutes in the deep-sea cold seep

The deep-sea samples were collected by *RV KEXUE* from a typical cold seep in the South China Sea (E119°17’07.322”, N22°06’58.598”) as described previously^21^. To understand the abundance of Tenericutes in deep-sea cold seep sediments, we selected and collected six sedimentary samples (RPC, ZC1, ZC2, ZC3, ZC4 and ZC5 at depth intervals of 0-10, 30-50, 90-110, 150-170, 210-230 and 230-250 cm, respectively) for operational taxonomic units (OTUs) sequencing performed by Novogene (Tianjin, China). Total DNAs from these samples were extracted by the CTAB/SDS method^28^ and were diluted to 1 ng/μL with sterile water and used for PCR template. 16S rRNA genes of distinct regions (16S V3/V4) were amplified using specific primers (341F: 5’-CCTAYGGGRBGCASCAG and 806R: 5’-GGACTACNNGGGTATCTAAT) with the barcode. These PCR products were purified with a Qiagen Gel Extraction Kit (Qiagen, Germany) and prepared to construct libraries. Sequencing libraries were generated using TruSeq® DNA PCR-Free Sample Preparation Kit (Illumina, USA) following the manufacturer’s instructions. The library quality was assessed on the Qubit@ 2.0 Fluorometer (Thermo Scientific) and Agilent Bioanalyzer 2100 system. And then the library was sequenced on an Illumina NovaSeq platform and 250 bp paired-end reads were generated. Paired-end reads were merged using FLASH (V1.2.7, http://ccb.jhu.edu/software/FLASH/)^29^, which was designed to merge paired-end reads when at least some of the reads overlap those generated from the opposite end of the same DNA fragments, and the splicing sequences were called raw tags. Quality filtering on the raw tags were performed under specific filtering conditions to obtain the high-quality clean tags^30^ according to the QIIME (V1.9.1, http://qiime.org/scripts/split_libraries_fastq.html) quality controlled process. The tags were compared with the reference database (Silva database, https://www.arb-silva.de/) using UCHIME algorithm (UCHIME Algorithm, http://www.drive5.com/usearch/manual/uchime_algo.html) to detect chimera sequences, and then the chimera sequences were removed^32^. And sequences analyses were performed by Uparse software (Uparse v7.0.1001, http://drive5.com/uparse/)^33^.

Sequences with ≥97% similarity were assigned to the same OTUs. The representative sequence for each OTU was screened for further annotation. For each representative sequence, the Silva Database (http://www.arb-silva.de/)^34^ was used based on Mothur algorithm to annotate taxonomic information.

### Metagenomic sequencing, assembly, binning and annotation

We selected three cold seep sediment samples (C1, C2 and C4, 20 g each) for metagenomic analysis in BGI (BGI, China). Briefly, total DNAs from these samples were extracted using the a Qiagen DNeasy® PowerSoil® Pro Kit (Qiagen, Germany) and the integrity of DNA was evaluated by gel electrophoresis. 0.5 μg DNA of each sample was used for libraries preparation with an amplification step. Thereafter, DNA was cleaved into 50 ~ 800 bp fragments using Covaris E220 ultrasonicator (Covaris, UK) and some fragments between 150 ~ 250 bp were selected using AMPure XP beads (Agencourt, USA) and repaired using T4 DNA polymerase (ENZYMATICS, USA). All NGS sequencing was performed on the BGISEQ-500 platform (BGI, China), generating 100 bp paired-end raw reads. Quality control was performed by SOAPnuke (v1.5.6) (setting: -l 20 -q 0.2 -n 0.05 -Q 2 -d -c 0 −5 0 −7 1)^35^ and the clean data were assembled using MEGAHIT (v1.1.3) (setting:--min-count 2 --k-min 33 --k-max 83 --k-step 10)^36^.

### Isolation and cultivation of deep-sea Izimaplasma

To enrich the Izimaplasma bacteria, the sediment samples were cultured at 28 °C for six months in an anaerobic enrichment medium containing (per liter seawater): 1.0 g NH_4_Cl, 1.0 g NaHCO_3_, 1.0 g CH_3_COONa, 0.5 g KH_2_PO_4_, 0.2 g MgSO_4_·7H_2_O, 1.0 mg *E. coli* genomic DNA, 0.7 g cysteine hydrochloride, 500 μL 0.1 % (w/v) resazurin and pH 7.0. This medium was prepared anaerobically as previously described^37^. During the course of enrichment, 1.0 mg genomic DNA was added once a month. After a six-month enrichment, 50 μL dilution portion was spread on Hungate tube covered by modified organic medium named ORG in this study, which contains 1 g yeast extract, 1 g peptone, 1 g NH_4_Cl, 1 g NaHCO_3_, 1 g CH_3_COONa, 0.5 g KH_2_PO_4_, 0.2 g MgSO_4_·7H_2_O, 0.7 g cysteine hydrochloride, 500 μL 0.1 % (w/v) resazurin, 1 L seawater, 15 g agar, pH7.0. The Hungate tubes were anaerobically incubated at 28 °C for 7 days. Individual colonies with distinct morphology were picked using sterilized bamboo sticks and then cultured in the ORG broth. Strain zrk13 was isolated and purified with ORG medium by repeated use of the Hungate roll-tube methods for several rounds until it was considered to be axenic. The purity of strain zrk13 was confirmed routinely by TEM and repeated partial sequencing of the 16S rRNA gene. Strain zrk13 is preserved at −80 °C in ORG broth supplemented with 20% (v/v) glycerol.

The temperature, pH and NaCl concentration ranges for the growth of strain zrk13 were determined in the ORG broth with incubation at 28 °C for 14 days. Growth assays were performed at different temperatures (4, 16, 28, 30, 37, 45, 60, 70, 80 °C). The pH range for growth was tested from pH 2.0 to pH 10.0 with increments of 0.5 pH units. Salt tolerance was tested in the modified ORG broth (with distilled water) supplemented with 0-10 % (w/v) NaCl (0.5 % intervals). Substrates utilization of strain zrk13 was tested in the medium (pH 7.0) consisting of (L^-1^): 5 g NaCl, 1 g NH_4_Cl, 0.5 g KH_2_PO4, 0.2 g MgSO_4_, 0.02 g yeast extract. Single substrate (including glucose, maltose, butyrate, fructose, sucrose, acetate, formate, starch, isomaltose, trehalose, galactose, cellulose, xylose, lactate, ethanol, D-mannose, glycerin, rhamnose and sorbitolurea) was added from sterile filtered stock solutions to the final concentration at 20 mM, respectively. Cell culture containing only 0.02 g yeast extract (L^-1^) without adding any other substrates was used as a control. All the cultures were incubated at 28 °C for 14 days. For each substrate, three biological replicates were performed.

### TEM observation

To observe the morphological characteristics of zrk13, its cell suspension was washed with Milli-Q water and centrifuged at 5,000 × *g* for 5 min, and taken by immersing copper grids coated with a carbon film for 20 min, washed for 10 min in distilled water and dried for 20 min at room temperature^38^. Ultrathin-section electron microscopic observation was performed as described previously^39,40^. Briefly, the samples were firstly preserved in 2.5% (v/v) glutaraldehyde overnight at 4 °C, washed three times with PBS and dehydrated in ethanol solutions of 30%, 50%, 70%, 90% and 100% for 10 min each time, and then the samples were embedded in a plastic resin. Ultrathin sections (50~70 nm) of cells were prepared with an ultramicrotome (Leica EM UC7), stained with uranyl acetate and lead citrate. All samples were examined using TEM (HT7700, Hitachi, Japan) with a JEOL JEM 12000 EX (equipped with a field emission gun) at 100 kV.

### Genome sequencing and genomic characteristics of strain zrk13

Genomic DNA was extracted from strain zrk13 cultured for 6 days at 28 °C. The DNA library was prepared using the Ligation Sequencing Kit (SQK-LSK109), and sequenced using a FLO-MIN106 vR9.4 flow-cell for 48 h on MinKNOWN software v1.4.2 (Oxford Nanopore Technologies (ONT), United Kingdom). Whole-genome sequence determinations of strain zrk13 were carried out with the Oxford Nanopore MinION (Oxford, United Kingdom) and Illumina MiSeq sequencing platform (San Diego, CA). A hybrid approach was utilized for genome assembly using reads from both platforms. Base-calling was performed using Albacore software v2.1.10 (Oxford Nanopore Technologies). Nanopore reads were processed using protocols toolkit for quality control and downstream analysis^41^. Filtered reads were assembled using Canu version 1.8^42^ using the default parameters for Nanopore data. And then the genome was assembled into a single contig and was manually circularized by deleting an overlapping end.

The genome relatedness values were calculated by multiple approaches: Average Nucleotide Identity (ANI) based on the MUMMER ultra-rapid aligning tool (ANIm), ANI based on the BLASTN algorithm (ANIb), the tetranucleotide signatures (Tetra), and *in silico* DNA-DNA similarity. ANIm, ANIb, and Tetra frequencies were calculated using JSpecies WS (http://jspecies.ribohost.com/jspeciesws/). The recommended species criterion cut-offs were used: 95% for the ANIb and ANIm and 0.99 for the Tetra signature. The amino acid identity (AAI) values were calculated by AAI-profiler (http://ekhidna2.biocenter.helsinki.fi/AAI/). The *in silico* DNA-DNA similarity values were calculated by the Genome-to-Genome Distance Calculator (GGDC) (http://ggdc.dsmz.de/)^43^. The *is*DDH results were based on the recommended formula 2, which is independent of genome size and, thus, is robust when using whole-genome sequences. The prediction of genes involved in DNA degradation for individual genomes was performed using Galaxy (Galaxy Version 2.6.0, https://galaxy.pasteur.fr/)^44^ with the NCBI BLAST+ blastp method.

### Phylogenetic analysis

The 16S rRNA gene tree was constructed with the full-length 16S rRNA sequences by the neighbor-joining algorithm, maximum likelihood and minimum-evolution methods. The full-length 16S rRNA gene sequence (1,537 bp) of strin zrk13 was obtained from the genome (accession number MW132883), and other related taxa used for phylogenetic analysis were obtained from NCBI (www.ncbi.nlm.nih.gov/). Phylogenetic trees were constructed using W-IQ-TREE web server (http://iqtree.cibiv.univie.ac.at)^45^ with LG+F+I+G4 model. The online tool Interactive Tree of Life (iTOL v4)^46^ was used for editing the tree.

### Growth assay of *X. coldseepsis* zrk13

Growth assays were performed at atmospheric pressure. Briefly, 16 mL strain zrk13 culture was inoculated in 2 L Hungate bottles containing 1.6 L of inorganic medium supplemented with or without 1.0 mg/L *E. coli* genomic DNA, ORG medium supplemented with 100 mM Na_2_S_2_O_3_ and oligotrophic medium (0.1 g/L yeast extract, 0.1 g/L peptone, 1.0 g/L NH_4_Cl, 1.0 g/L NaHCO_3_, 1.0 g/L C¾COONa, 0.5 g/L KH_2_PO_4_, 0.2 g/L MgSO_4_·7H_2_O, 0.7 g/L cysteine hydrochloride, 500 μL/L of 0.1 % (w/v) resazurin and pH 7.0), respectively. These Hungate bottles were anaerobically incubated at 28 °C. Bacterial growth status was monitored by measuring the OD600 value every day until cell growth reached the stationary phase. Since strain zrk13 grew very slow in the inorganic medium, we also adopted the quantitative PCR (qPCR) to measure its growth status.

### Detection of DNA degradation by strain zrk13

Electrophoresis was performed to detect the ability of DNA degradation of strain zrk13. Briefly, the *E. coli* genomic DNA and RNA were extracted from overnight cultured *E. coli* DH5α cells with a Genomic DNA and RNA Kit (Tsingke, China) by omiting the RNA removal step. The reaction system (20 μL) contained 2 μL bacterial supernatant, 2 μL buffer and 16 μL *E. coli* genomic DNA and rRNA extracts (2 μg). In parallel, the supernatant of a Clostridia bacterium was used as a control. All bacterial supernatants were obtained by centrifugation with 12,000 *g* for 10 min from 10-day cultures. Assays were performed at 37 °C for 5 min, 10 min and 15 min, respectively. Finally, the reaction solutions were detected by 1% agarose gel electrophoresis at 180 V for 20 min. The imaging of the gel was taken by the Gel Image System (Tanon 2500, China).

### Quantitative real-time PCR assay

For qRT-PCR, cells of strain zrk13 were cultured in inorganic medium supplemented with or without 1.0 mg/L *E. coli* genomic DNA or organic medium at 28 °C for 15 days with harvesting cells from 100 mL medium every day. Total RNAs from each sample were extracted using the Trizol reagent (Solarbio, China) and the RNA concentration was measured using Qubit® RNA Assay Kit in Qubit® 2.0 Flurometer (Life Technologies, CA, USA). Then RNA from corresponding sample was reverse transcribed into cDNA and the transcriptional levels of different genes were determined by qRT-PCR using SybrGreen Premix Low rox (MDbio, China) and the QuantStudioTM 6 Flex (Thermo Fisher Scientific, USA). The PCR condition was set as following: initial denaturation at 95 °C for 3 min, followed by 40 cycles of denaturation at 95 °C for 10 s, annealing at 60 °C for 30 s, and extension at 72 °C for 30 s. 16S rRNA was used as an internal reference and the gene expression was calculated using the 2^-ΔΔCt^ method, with each transcript signal normalized to that of 16S rRNA. Transcript signals for each treatment were compared to those of control group. Specific primers for genes related to DNA degradation and 16S rRNA were designed using Primer 5.0 as shown in Supplementary Table 4. The standard curve was generated using five independent dilution series of plasmid DNA carrying a fragment (143 bp) of the strain zrk13^T^ 16S rRNA gene. This qPCR assay standard curve had a slope of −2.5162, a y-intercept of 30.53 and an R^2^ value of 0.9924. All qRT-PCR runs were performed in three biological and three technical replicates.

### Transcriptomic analysis

Transcriptomic analysis was performed by Novogene (Tianjin, China). Briefly, cells suspension of *X. coldseepsis* zrk13 cultured in oligotrophic medium, ORG medium supplemented with or without 100 mM Na2S2O3 for 7 days was respectively collected for further transcriptomic analysis. All cultures were performed in 2 L anaerobic bottles. For *in situ* experiments, *X*. *coldseepsis* zrk13 was firstly cultured in ORG medium for 5 days, and then was divided into two parts: one part was divided into three gas samples bags (which not allowing any exchanges between inside and outside; Aluminum-plastic composite film, Hede, China) and used as the control group; the other part was divided into three dialysis bags (8,000-14,000 Da cutoff, which allowing the exchanges of substances smaller than 8,000 Da but preventing bacterial cells from entering or leaving the bag; Solarbio, China) and used as the experimental group. All the samples were placed simultaneously in the deep-sea cold seep (E119°17’04.429”, N22°07’01.523”) for 10 days in the June 2020 during the cruise of *Kexue* vessel. After 10 days incubation in the deep sea, the bags were taken out and the cells were immediately collected and saved in the −80 freezer until further use. Thereafter, the cells were checked by 16S rRNA sequencing to confirm the purity of the culture and performed further transcriptomic analysis. For transcriptomic analyses, total *X*. *coldseepsis* zrk13 RNAs were extracted using TRIzol reagent (Invitrogen, USA) and DNA contamination was removed using the MEGA clear™ Kit (Life technologies, USA). Detailed protocols of the following procedures including library preparation, clustering and sequencing and data analyses were described in the Supplementary Information.

## Data availability

The whole 16S rRNA sequence of *X. coldseepsis* zrk13 has been deposited in the GenBank database (accession number MW132883). The complete genome sequence of *X*. *coldseepsis* zrk13 has been deposited at GenBank under the accession number CP048914. The raw sequencing reads for transcriptomic analysis have been deposited to NCBI Short Read Archive (accession numbers: PRJNA664657, PRJNA669477 and PRJNA669478). The raw amplicon sequencing data have also been deposited to NCBI Short Read Archive (accession number: PRJNA675395).

## Acknowledgements

This work was funded by the Strategic Priority Research Program of the Chinese Academy of Sciences (Grant No. XDA22050301), China Ocean Mineral Resources R&D Association Grant (Grant No. DY135-B2-14), National Key R and D Program of China (Grant No. 2018YFC0310800), the Taishan Young Scholar Program of Shandong Province (tsqn20161051), and Qingdao Innovation Leadership Program (Grant No. 18-1-2-7-zhc) for Chaomin Sun. This study is also funded by the Open Research Project of National Major Science & Technology Infrastructure *(RV KEXUE)* (Grant No. NMSTI-KEXUE2017K01).

## Author contributions

RZ and CS conceived and designed the study; RZ conducted most of the experiments; RL, YS and GL collected the samples from the deep-sea cold seep; RC helped to analyze the metagenomes; RZ and CS lead the writing of the manuscript; all authors contributed to and reviewed the manuscript.

## Supplementary methods

### Transcriptional profiling of *X. coldseepsis* zrk13 cultured in different conditions

#### (1) Library preparation for strand-specific transcriptome sequencing

A total amount of 3 μg RNA per sample was used as input material for the RNA sample preparations. Sequencing libraries were generated using NEBNext® Ultra™ Directional RNA Library Prep Kit for Illumina® (NEB, USA) following manufacturer’s recommendations and index codes were added to attribute sequences to each sample. rRNA is removed using a specialized kit that leaves the mRNA. Fragmentation was carried out using divalent cations under elevated temperature in NEBNext First Strand Synthesis Reaction Buffer (5×). First strand cDNA was synthesized using random hexamer primer and M-MuLV Reverse Transcriptase (RNaseH^-^). Second strand cDNA synthesis was subsequently performed using DNA Polymerase I and RNase H. In the reaction buffer, dNTPs with dTTP were replaced by dUTP. Remaining overhangs were converted into blunt ends via exonuclease/polymerase activities. After adenylation of 3’ ends of DNA fragments, NEBNext Adaptor with hairpin loop structure was ligated to prepare for hybridization. In order to select cDNA fragments of preferentially 150~200 bp in length, the library fragments were purified with AMPure XP system (Beckman Coulter, Beverly, USA). Then 3 μL USER Enzyme (NEB, USA) was used with size-selected, adaptor-ligated cDNA at 37 °C for 15 min followed by 5 min at 95 °C before PCR. Then PCR was performed with Phusion High-Fidelity DNA polymerase, Universal PCR primers and Index (X) Primer. At last, products were purified (AMPure XP system) and library quality was assessed on the Agilent Bioanalyzer 2100 system.

#### (2) Clustering and sequencing

The clustering of the index-coded samples was performed on a cBot Cluster Generation System using TruSeq PE Cluster Kit v3-cBot-HS (Illumia) according to the manufacturer’s instructions. After cluster generation, the library preparations were sequenced on an Illumina Hiseq platform and paired-end reads were generated.

#### (3) Data analysis

Raw data (raw reads) of fastq format were firstly processed through in-house perl scripts. In this step, clean data (clean reads) were obtained by removing reads containing adapter, reads containing ploy-N and low quality reads from raw data. At the same time, Q20, Q30 and GC content the clean data were calculated. All the downstream analyses were based on the clean data with high quality. Reference genome and gene model annotation files were downloaded from genome website directly. Both building index of reference genome and aligning clean reads to reference genome were used Bowtie2-2.2.3^1^. HTSeq v0.6.1 was used to count the reads numbers mapped to each gene. And then FPKM of each gene was calculated based on the length of the gene and reads count mapped to this gene. FPKM, expected number of Fragments Per Kilobase of transcript sequence per Millions base pairs sequenced, considers the effect of sequencing depth and gene length for the reads count at the same time, and is currently the most commonly used method for estimating gene expression levels^2^.

#### (4) Differential expression analysis

Differential expression analysis of two conditions/groups (two biological replicates per condition) was performed using the DESeq R package (1.18.0)^3^. DESeq provide statistical routines for determining differential expression in digital gene expression data using a model based on the negative binomial distribution. The resulting *P*-values were adjusted using the Benjamini and Hochberg’s approach for controlling the false discovery rate. Genes with an adjusted *P*-value < 0.05 found by DESeq were assigned as differentially expressed. (For DEGSeq without biological replicates) Prior to differential gene expression analysis, for each sequenced library, the read counts were adjusted by edgeR program package through one scaling normalized factor. Differential expression analysis of two conditions was performed using the DEGSeq R package (1.20.0)^4^. The *P* values were adjusted using the Benjamini & Hochberg method. Corrected *P*-value of 0.005 and log_2_ (Fold change) of 1 were set as the threshold for significantly differential expression.

#### (5) GO and KEGG enrichment analysis of differentially expressed genes

Gene Ontology (GO) enrichment analysis of differentially expressed genes was implemented by the GOseq R package, in which gene length bias was corrected^5^. GO terms with corrected *P* value less than 0.05 were considered significantly enriched by differential expressed genes. KEGG is a database resource for understanding high-level functions and utilities of the biological system, such as the cell, the organism and the ecosystem, from molecular-level information, especially large-scale molecular datasets generated by genome sequencing and other high-throughput experimental technologies (http://www.genome.jp/kegg/)^6^. We used KOBAS software to test the statistical enrichment of differential expression genes in KEGG pathways.

## Supplementary results

**Supplementary Fig 1.**
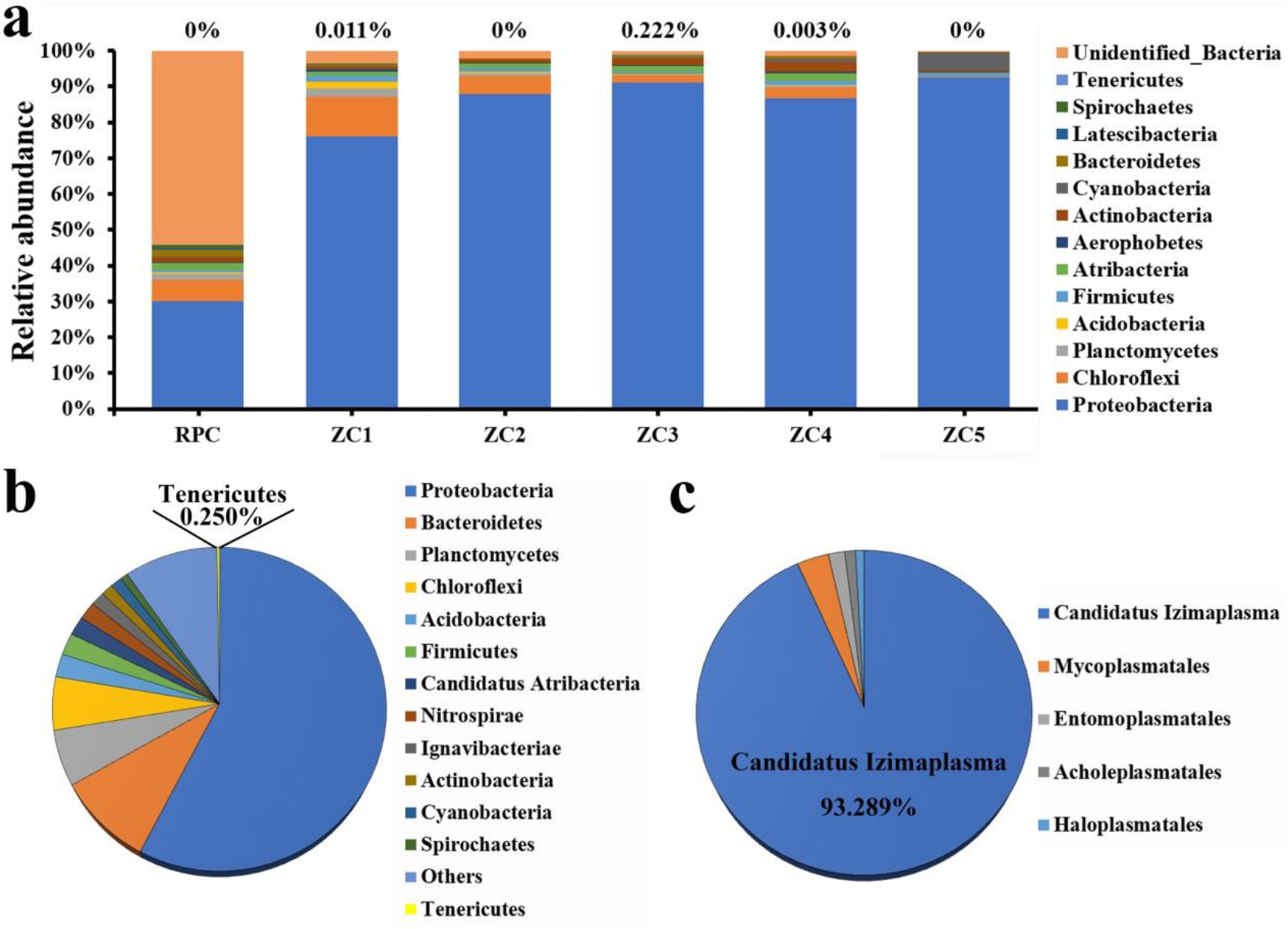
Quantification of the abundance of Tenericutes and *Candidatus* Izimaplasma in the deep-sea cold seep. **a**, The community structure of six sampling sites in the cold seep sediments as revealed by 16S rRNA gene amplicon profiling. The relative abundances of operational taxonomic units (OTUs) representing different bacteria are shown at the phylum level. **b**, Quantification of the abundance of Tenericutes in the bacteria domain based on the metagenomics sequencing. **c**, Quantification of the abundance of *Candidatus* Izimaplasma in the phylum Tenericutes based on the metagenomics sequencing.

**Supplementary Fig 2.**
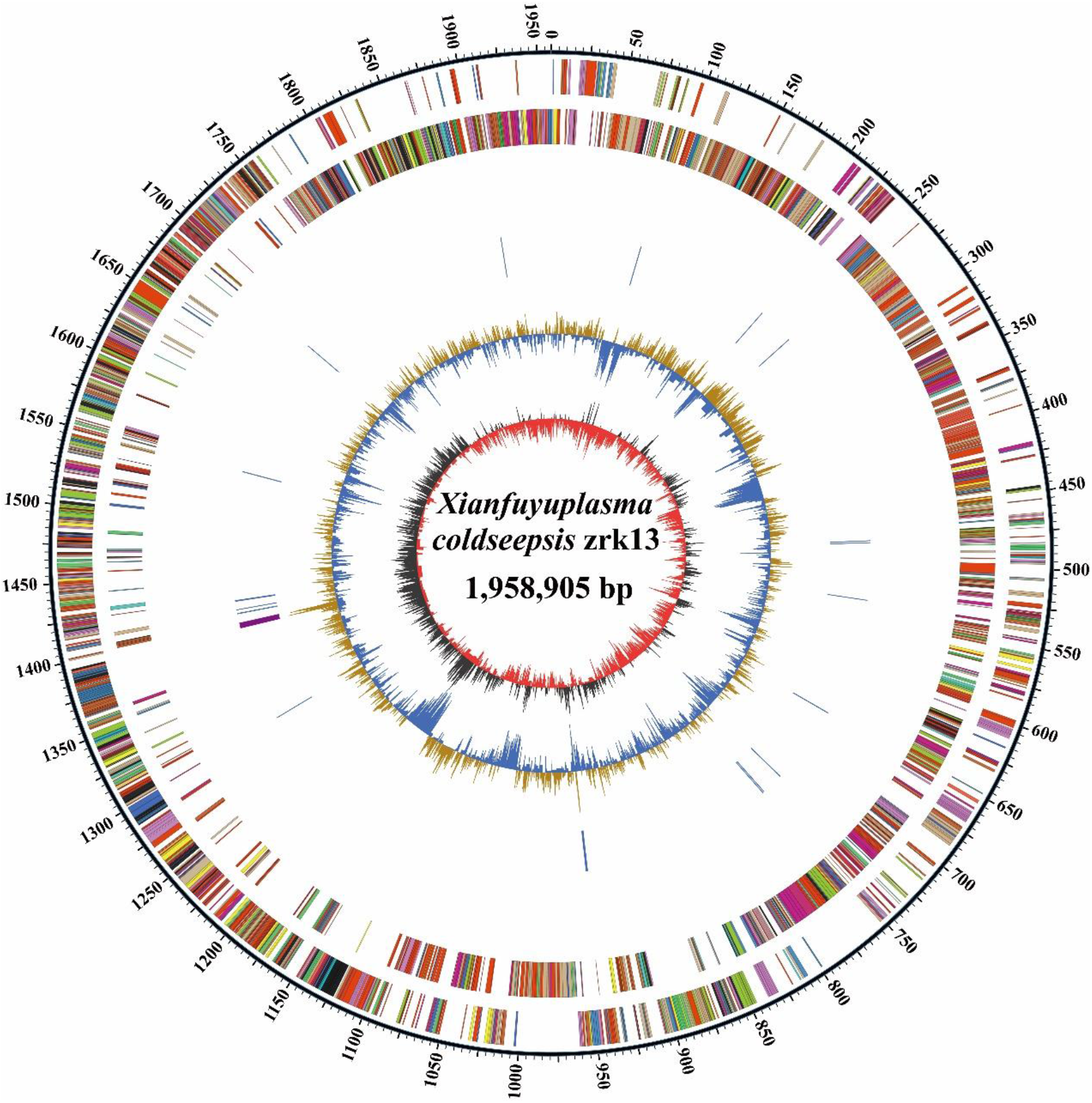
Circular diagram of the genome of *X.coldseepsis* zrk13. Rings indicate, from outside to the center: a genome-wide marker with a scale of 50 kb; forward strand genes, colored by COG category; reverse strand genes, colored by COG category; repetitive sequences; RNA genes (tRNAs blue, rRNAs purple); GC content; GC skew.

**Supplementary Fig 3.**
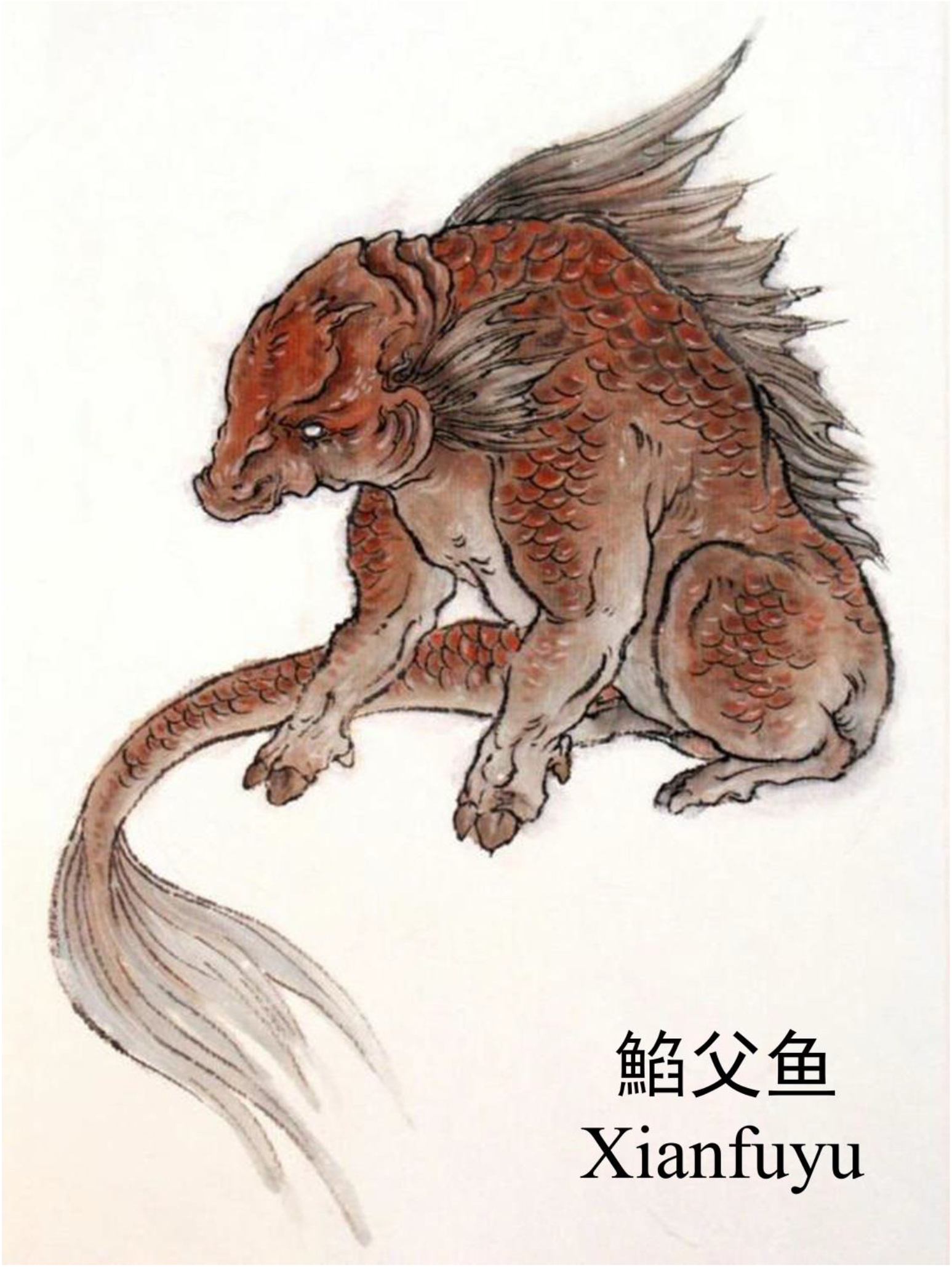
The picture of ‘Xianfuyu’ comes from a strange animal recorded in a very famous Chinese mythology-Classic of Mountains and Rivers. The ‘Xianfuyu’ is the father of all the fishes, which looks like gouramies, with a fish head and a pig body.

**Supplementary Fig 4.**
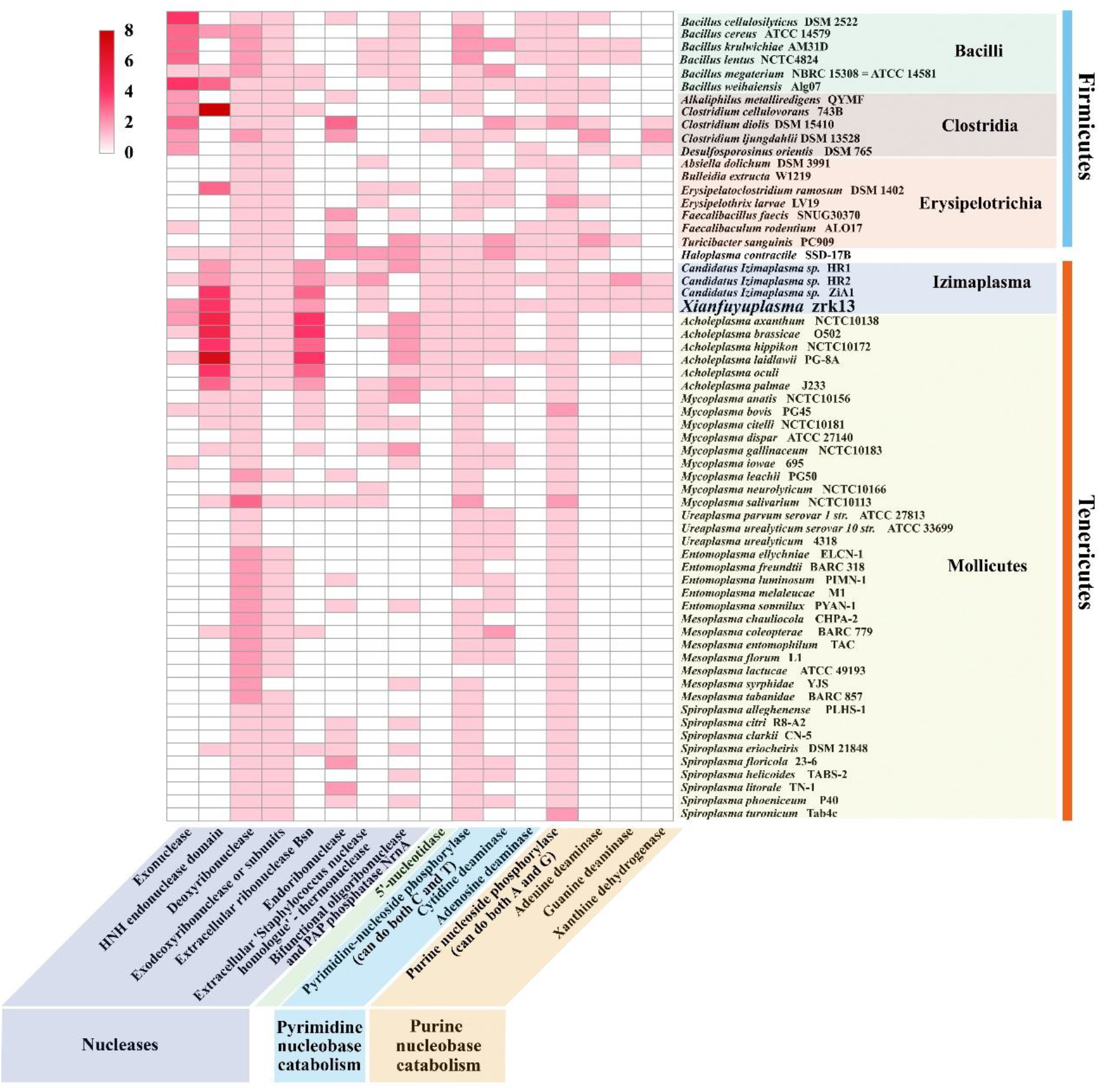
Distribution of key genes associated with DNA degradation in the the genomes of typical species of phylums Tenericutes and Firmicutes.

**Supplementary Table 1.**
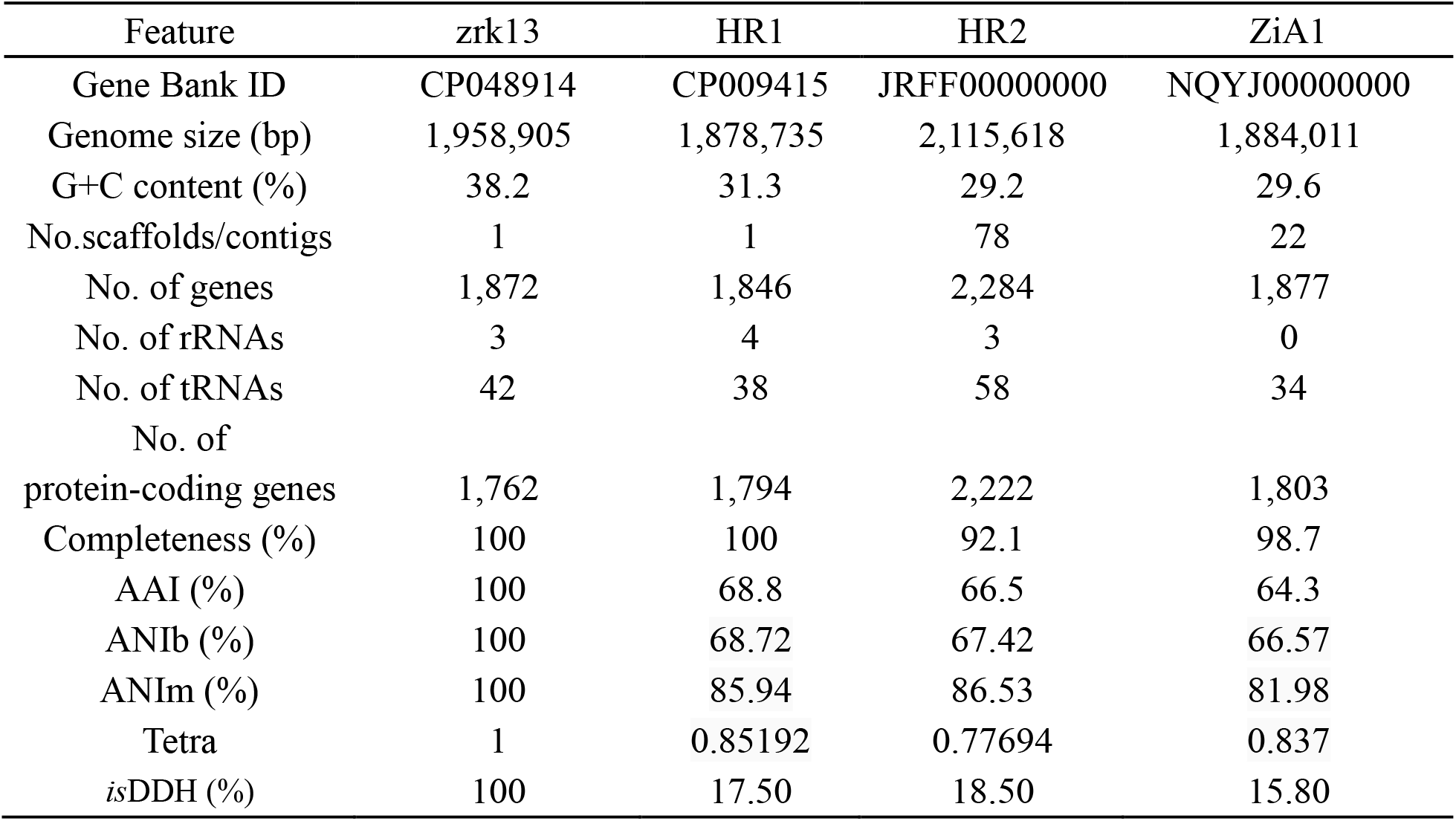
Genomic features of *X. coldseepsis* zrk13 with other uncultivated Izimaplasma strains HR1, HR2 and ZiA1.

**Supplementary Table 2.** Physiological characteristics of *X. coldseepsis* zrk13.

| Characteristic | zrk13 |
| --- | --- |
| Cell length (µm) | 0.2-0.5 |
| Temperature range for growth ( °C) | 24-32 |
| Optimum | 28 |
| pH range for growth | 6.0-8.0 |
| Optimum | 7.0 |
| NaCl range for growth (%) | 0-8 |
| Optimum | 4 |
| Utilization as a sole carbon source: |  |
| Glucose | + |
| Maltose | + |
| Butyrate | + |
| Fructose | + |
| Sucrose | + |
| Acetate | + |
| Formate | + |
| Starch | + |
| Isomaltose | + |
| Trehalose | + |
| Galactose | – |
| Cellulose | – |
| Xylose | – |
| Lactate | + |
| Ethanol | + |
| D-mannose | + |
| Glycerin | + |
| Rhamnose | + |
| Sorbitol | – |
| DNA G+C content (mol%) | 38.21 % |
| Isolation source | deep-sea sediments |

